# Nerve recovery from treatment with a vascularized nerve graft compared to an autologous non-vascularized nerve graft in animal models: a systematic review and meta-analysis

**DOI:** 10.1101/2021.05.13.443972

**Authors:** Berend O. Broeren, Liron S. Duraku, Caroline A. Hundepool, Erik T. Walbeehm, J. Michiel Zuidam, Carlijn R. Hooijmans, Tim De Jong

## Abstract

**Background:** Treatment of nerve injuries proves to be a worldwide clinical challenge. Vascularized nerve grafts are suggested to be a promising alternative for bridging a nerve gap to the current gold standard, an autologous non-vascularized nerve graft. However, there is no adequate clinical evidence for the beneficial effect of vascularized nerve grafts and they are still disputed in clinical practice.

**Objective:** To systematically review whether vascularized nerve grafts give a superior nerve recovery compared to non-vascularized nerve autografts regarding histological and electrophysiological outcomes in animal models.

**Material and methods:** PubMed and Embase were systematically searched. The inclusion criteria were as follows: 1) the study was an original full paper which presented unique data; 2) a clear comparison between a vascularized and a non-vascularized autologous nerve transfer was made; 3) the population study were animals of all genders and ages. A standardized mean difference and 95% confidence intervals for each comparison was calculated to estimate the overall effect. Subgroup analyses were conducted on graft length, species and time frames.

**Results:** Fourteen articles were included in this review and all of them were included in the meta-analyses. A vascularized nerve graft resulted in a significantly larger diameter, higher nerve conduction velocity and axonal count compared to an autologous non-vascularized nerve

graft. However, during sensitivity analysis the effect on axonal count disappeared. No significant difference was observed in muscle weight.

**Conclusion:** Treating a nerve gap with a vascularized graft results in superior nerve recovery compared to non-vascularized nerve autografts in three out of four outcome measurements. However, this conclusion needs to be taken with some caution due to the inherent limitations of this meta-analysis. We recommend future studies to be performed under conditions more closely resembling human circumstances and to use long nerve defects.

## Introduction

Treatment of nerve injuries proves to be a worldwide clinical challenge. Even though adequately treated, affected patients may suffer from chronic pain or lasting motor and sensory deficits.(1) For clinical situations in which it is necessary to bridge a nerve gap, the current gold standard is an autologous non-vascularized (conventional) nerve graft. A nerve graft always has a worse outcome compared to primary coaptation, due to two anastomosis sides, ischemia of the graft and frequently a poor wound bed.(2)

To improve the outcome after nerve repair with conventional nerve autografts the blood supply can be taken along with the nerve graft, the so-called vascularized nerve graft. Grafted nerves need considerable energy to regenerate and to maintain function. This energy is delivered by the intraneural vascular system, which is connected to extrinsic vessels. Therefore, an instant and sufficient blood supply may be beneficial for recovery.(3),(4),(5)

There is no adequate clinical evidence of beneficial effect of vascularized nerve grafts except several case reports and case series.(6),(7),(8),(9),(10),(11),(12) The use of a vascularized nerve graft was first reported in 1976 by Taylor and Ham. They used 24 cm of the superficial radial nerve attached to the radial artery to reconstruct a median nerve. (13)

Since the first publication by Taylor and Ham, many experimental studies in animal models have been reported. Vascularized nerve grafts have been successfully attempted in rats, rabbits, dogs, and other species to develop a model that is feasible, straightforward, reliable, and reproducible.(14)

Nowadays, the use of vascularized nerve grafts is still debated in clinical practice because of several reasons: 1) the concern of a more significant donor site morbidity compared to conventional nerve autografts; 2) the lack of clinical evidence indicating the superiority of a vascularized nerve graft; 3) the difficulty to set up a controlled trial, due to the high heterogeneity of patients as well as nerve defects.

Therefore, a systematic review and meta-analysis of animal models was conducted to investigate whether vascularized nerve grafts show a superior nerve recovery compared to non-vascularized nerve autografts regarding histological and electrophysiological factors.

## Material and methods

### Research protocol

This systematic review protocol was defined in advance and registered in an international database (PROSPERO, registration number CRD42020184363).

### Search strategy

A systematic search has been performed in the PubMed (Medline) and Embase (OVID) databases to identify all original articles. The search included studies up to 26th of May 2020. Search terms included ‘nerve transfer’, ‘nerve graft’, ‘vascularized’ and ‘vascularization’ and their synonyms in abstract and title fields (for the complete search strategy, see S1 Table). The SYRCLE search filters to identify all animal studies were used.(15, 16) Duplicates were taken out using Endnote (Clarivate Analytics, Pennsylvania, USA). Two authors (BOB and TDJ) independently screened all titles and abstracts for their relevance utilizing predetermined inclusion and exclusion criteria. A reference- and citation check of the remaining studies was conducted manually to acquire potentially missed relevant articles. Afterward, the full text of the relevant articles was screened for final selection. Contradictory judgments were resolved by consensus discussion. No language or date restrictions were applied.

### Inclusion and exclusion criteria

Articles were included when 1) the study was an original full paper which presented unique data; 2) a clear comparison between a vascularized and a non-vascularized autologous nerve transfer was made; 3) the population study were animals (all species) of all genders and ages; 4) the study investigated the effects of vascularized nerve grafts on: axonal count, diameter, nerve conduction velocity and muscle weight. No language or publication date restrictions were applied.

### Critical appraisal

All included studies were appraised using the SYRCLE’s tool for assessing the risk of bias for animal studies.(17) This appraisal was done by two authors (BOB and TDJ) independently and subsequently merged by consensus. All criteria were scored a “yes” indicating a low risk of bias or a “no” indicating high risk of bias or a “?” indicating an unknown risk of bias. Baseline characteristics were: weight, age and race. Selective outcome reporting was determined by establishing if all outcome measures mentioned in material and methods were reported in the results section as well. To compensate for judging a lot of items as “unclear risk of bias” due to highly inadequate reporting of experimental details on animals, methods and materials, we included two items. The first item was reporting on any measure of randomization and the second item was reporting on any measure of blinding. Here a “yes” signifies reported and a “no” means unreported.

### Data extraction

Data were in duplicate extracted from the selected studies by two authors (BOB and TDJ). The descriptive data included: publication year, first author’s name, studied species, gender, total number of animals, total grafts, studied nerve, studied muscle, graft length and time points. For the meta-analysis, the mean, sd and n of the following outcomes were extracted for axonal count, diameter, nerve conduction velocity and muscle weight. When measurements of multiple locations per nerve were reported, the most distal segment of the graft was used. In case the SEM was reported it was converted to SD (SD = SEM x √n). When outcome measure data was missing, authors were contacted for additional information. When data were displayed only graphically, we used Universal Desktop Ruler software (https://avpsoft.com/products/udruler/), to determine an adequate estimation of the outcome measurements. The mean of two independent measurements was used.

### Statistical analysis

Data were analyzed using Review Manager, Version 5.4. Copenhagen: The Nordic Cochrane Centre, The Cochrane Collaboration. Meta-analysis was performed for all four outcome measurements by calculating the standardized mean difference (SMD) between vascularized and conventional grafts. Whenever a comparison reported an SD of 0 it was excluded from meta-analysis. A random effects model was applied, taking into account the accuracy of independent studies and the variation among studies and weighing all studies accordingly. Heterogeneity was measured using I^2^. Subgroup analyses were performed for different species (rabbit and rat), different graft length (0-2 cm, 2-4 cm and 4 > cm) and different time frames (0-2 months, 2-4 months and 4 > months). The results of subgroup analysis were only interpreted when groups consisted of 3 or more individual studies.

Funnel plots, egger regression and Trim and Fill analysis were used to search for evidence for publication bias if at least 10 or more studies per outcome. Because SMDs may cause funnel plot distortion, we plotted the SMD against a sample size-based precision estimate(1/√(n)).

To assess the robustness of our findings, a sensitivity analysis was performed. We evaluated the impact of excluding studies which used animals as their own control group.

## Results

### Study selection process

The search strategy presented in S1 Table retrieved 303 records, including 131 in PubMed and 172 in Embase. After removing duplicates, 203 articles appeared to be unique (Fig 1. shows a consort flow chart). After title abstract screening, 28 studies entered the full text screening phase. Finally, 14 articles were included in the review.

**Fig 1.**
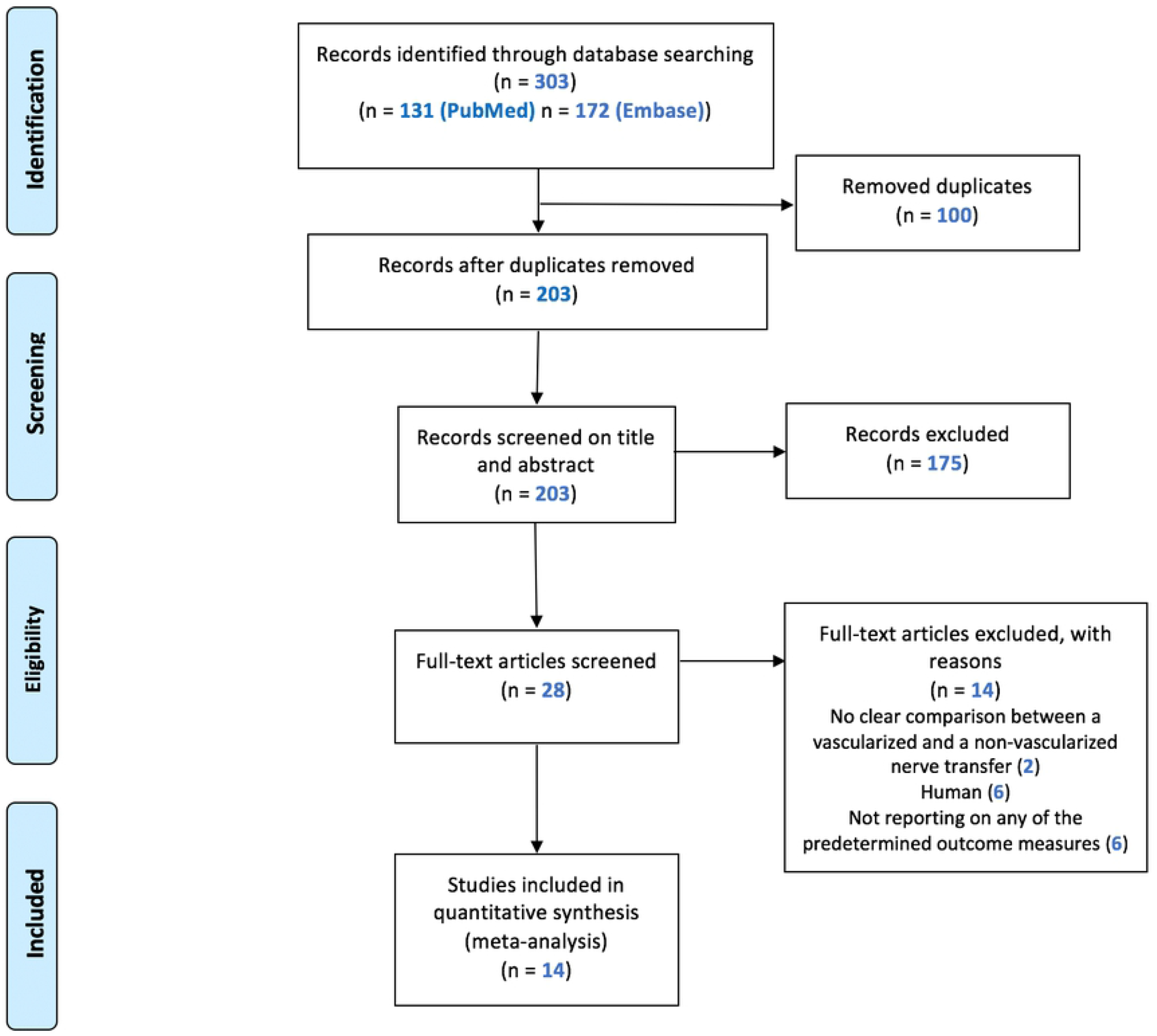
Flow chat of the study selection.

### Study quality and risk of bias

This review clearly revealed that methodological details of animal experiments were often poorly reported. Reporting about any randomization and blinding measures taken in the conducted studies was respectively 21% (3 out of 14 publications)

The general results of our risk of bias assessment of the included references in this review are presented in Fig 2. Poor reporting of essential methodological details in most animal experiments resulted in an unclear risk of bias in the majority of studies. Risk of bias was scored separately for the 3 studies that used animals as their own control group because some aspects were not applicable (Fig 3).

**Fig 2.**
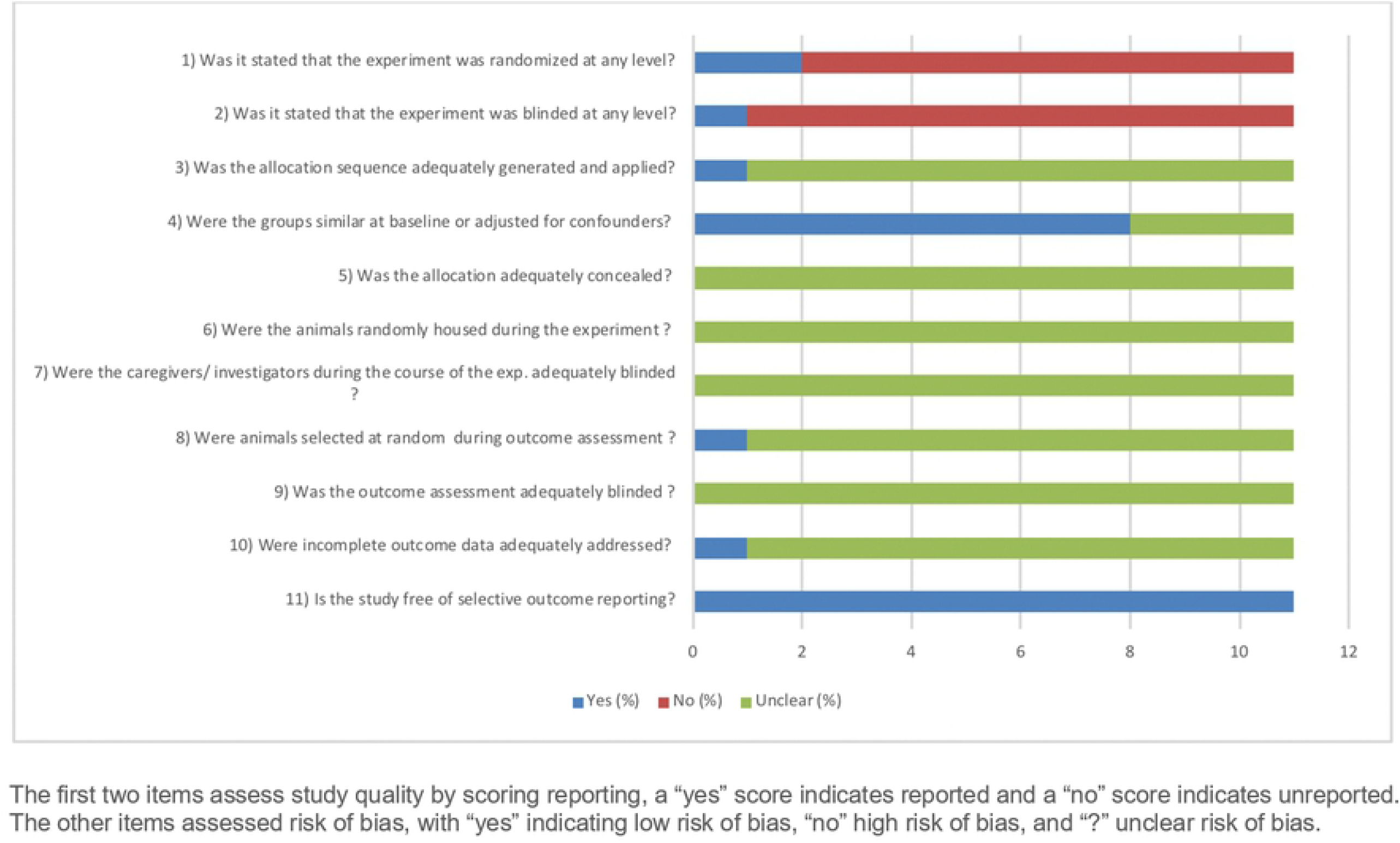
Result of the risk of bias assessment of 11 included studie.

**Fig 3.**
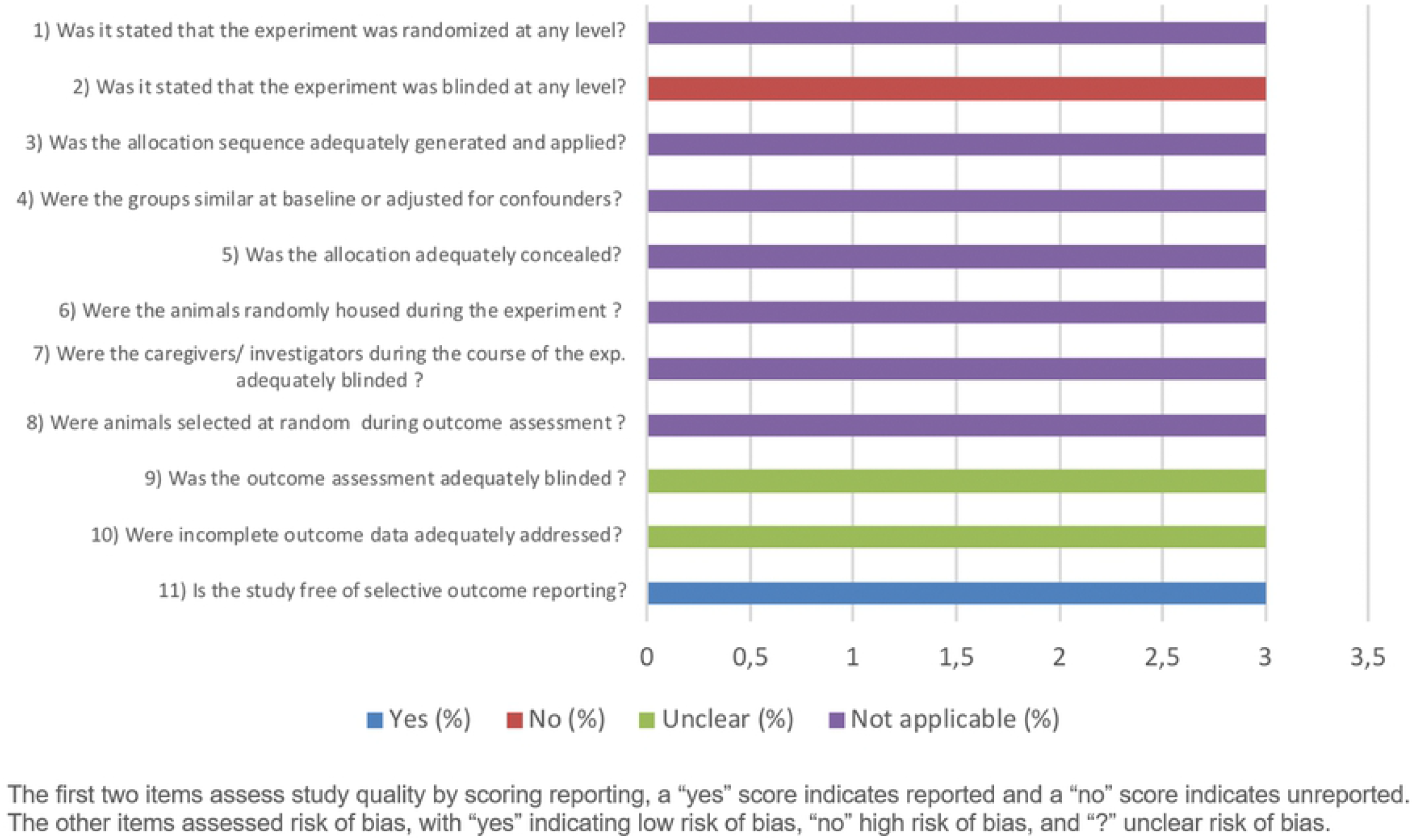
Result of the risk of bias assessment of 3 included studie.

### Study characteristics

The characteristics of the 14 included publications are shown in Table 1.(18-31) All studies used either a rabbit (57%) or rat (43%) model. Notably, more than half the studies did not report gender (8 out of 14 studies). Out of the remaining studies 3 used females, 2 used males and in one both females and males were used. The sciatic nerve was the most commonly used nerve (50%), followed by the median nerve (29%), facial nerve (7%), peroneal nerve (7%) and auricular nerve (7%).

**Table 1.** The characteristics of all 14 included references.

| *Reference* | *Outcome measurements* | *Species* | *Gender* | *Animals* | *Grafts* | *Nerve* | *Muscle* | *Graft size (mm)* | *Time points (days)* |
| --- | --- | --- | --- | --- | --- | --- | --- | --- | --- |
| Bertelli et al., 1996 | Muscle weight | Rat | Female | 70 | 70 | Median | FCR | 20 | 95, 120, 150, 210, 360 |
| Donzelli et al., 2016 | Axonal count<br>Diameter | Rabbit | Male | 20 | 20 | Sciatic |  |  | 30, 90 |
| Hems et al., 1992 | Axonal count | Rabbit | NR | 8 | 8 | Peroneal |  | 50 | 250 |
| Kanaya et al., 1992 | Axonal count<br>Nerve conduction velocity<br>Muscle weight | Rat | Female | 22 | 22 | Sciatic | Tibialis anterior | 25 | 84 |
| Kawai et al., 1990 | Axonal count<br>Diameter | Rabbit | NR | 34 | 67 | Median |  | 20,40, 60 | 56, 168 |
| Koshima et al., 1985 | Axonal count<br>Diameter | Rat | Male | 38 | 38 | Sciatic |  | 15 | 28, 56, 84, 112, 140, 168 |
| Koshima.2 et al., 1985 | Axonal count<br>Diameter | Rat | NR | 74 | 74 | Sciatic |  | 15 | 21, 28, 35, 42, 49, 56, 84, 112, 140, 168, 224 |
| Mani et al., 1992 | Diameter | Rabbit | Male/<br>Female | 11 | 11 | Sciatic |  | 30 | 308 |
| Matsumine et al., 2013 | Axonal count<br>Diameter | Rat | NR | 14 | 14 | Median |  | 7 | 210 |
| Ozcan et al., 1993 | Axonal count<br>Diameter | Rabbit | Female | 10 | 10 | facial |  | 10 | 84 |
| Seckel et al., 1986 | Axonal count | Rat | NR | 13 | 26 | Sciatic |  | 10 | 21, 28, 42 |
| Shibata et al., 1988 | Axonal count<br>Diameter<br>Nerve conduction velocity | Rabbit | NR | 39 | 39 | Median |  | 30 | 70, 168 |
| Tark et al., 2001 | Axonal count | Rabbit | NR | 33 | 66 | Sciatic |  | 40 | 56, 84, 112 |
| Zhu et al., 2015 | Nerve conduction velocity | Rabbit | NR | 6 | 6 | Auricular |  | 20 | 112 |

NR: not reported
FCR: flexor carpi radialis

### Axonal count

Data on axonal count could be retrieved from 11 independent studies containing 37 comparisons.(19-24, 26-30) Seven comparisons had to be excluded because not all outcome data was available. Out of the remaining 30 experiments conducted, data obtained from rabbits and rats was both 50%. In total 352 grafts were placed in 309 animals.

There was a variation in graft length from 7 to 60 mm. The graft length was unreported in two of the comparisons. Data were extracted at different time points varying between 21 and 250 days.

Overall analysis showed a significant difference in favor of treatment with a vascularized nerve graft (SMD, 0.46 [95% CI 0.06 to 0.86], N = 30) (Fig 4). The overall between study heterogeneity was moderate to high at I^2^ = 61%.

**Fig 4.**
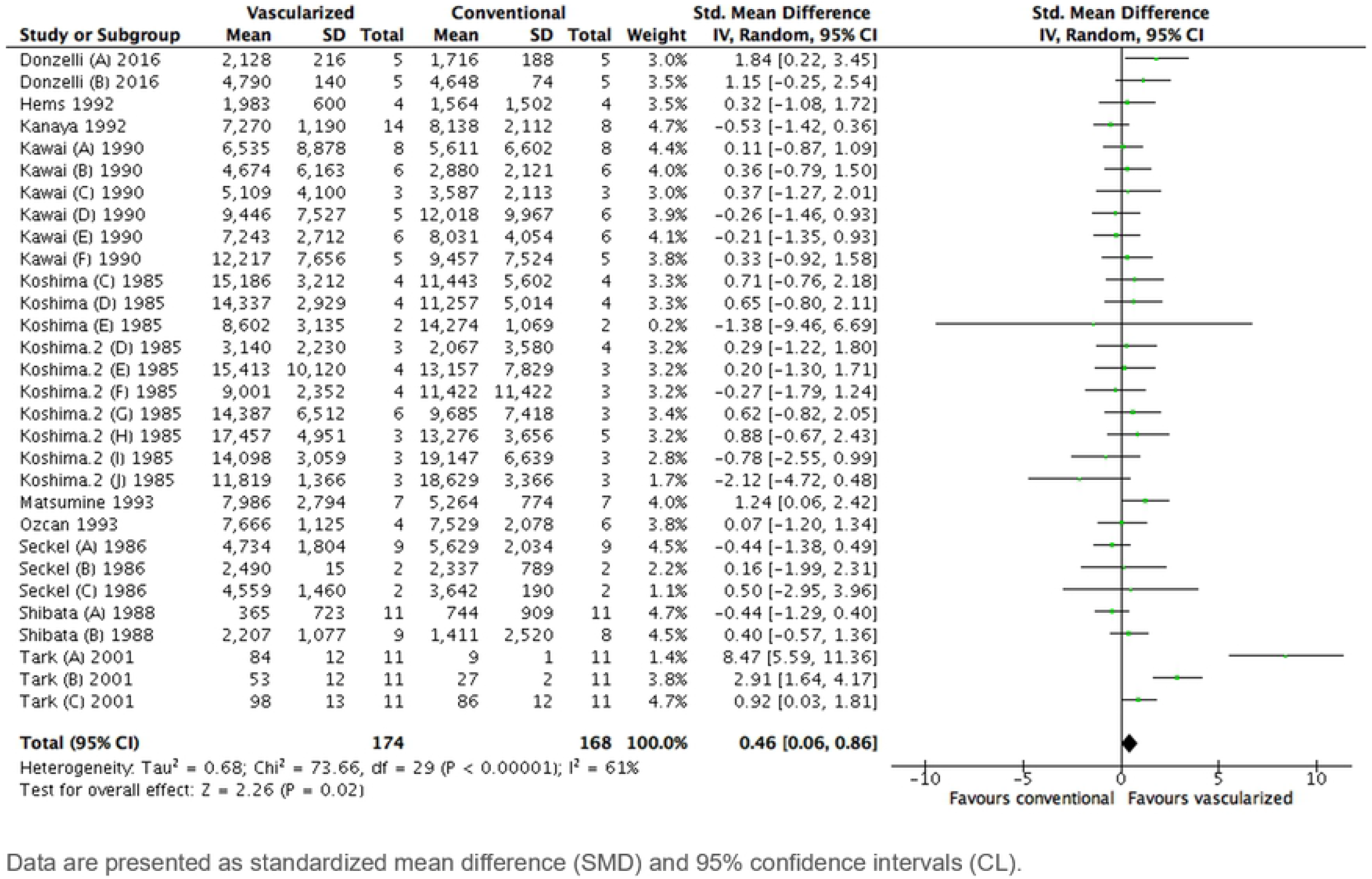
Forest plot of the effect of treatment with a vascularized n.

Subgroup analyses revealed no differences in graft length, species and time frames when comparing axonal count between vascularized and conventional nerve autografts. The graft length middle group consisted of too few studies for subgroup analyses. (S1 Fig, S2 Fig, S3 Fig).

### Diameter

Eight studies, containing 31 comparisons, reported nerve fiber diameter on histological examination.(19, 22-27, 29) Since not all data was available, 10 of the 31 comparisons had to be excluded. Rabbits and rats were used in 52% and 48% respectively. All 21 comparisons combined, a total of 148 animals were operated on, resulting in 185 grafts that met our selection criteria. Graft length varied between 7 and 60 mm. In one of the studies, it was unclear which graft length was used. The time points at which data were extracted ranged from 21 to 308 days.

Analysis of all 21 included comparisons showed a significantly larger diameter after treatment with a vascularized nerve graft (SMD, 0.59 [95% CI 0.16 to 1.02], N = 21) (Fig 5). Study heterogeneity was moderate (I^2^ = 36%).

**Fig 5.**
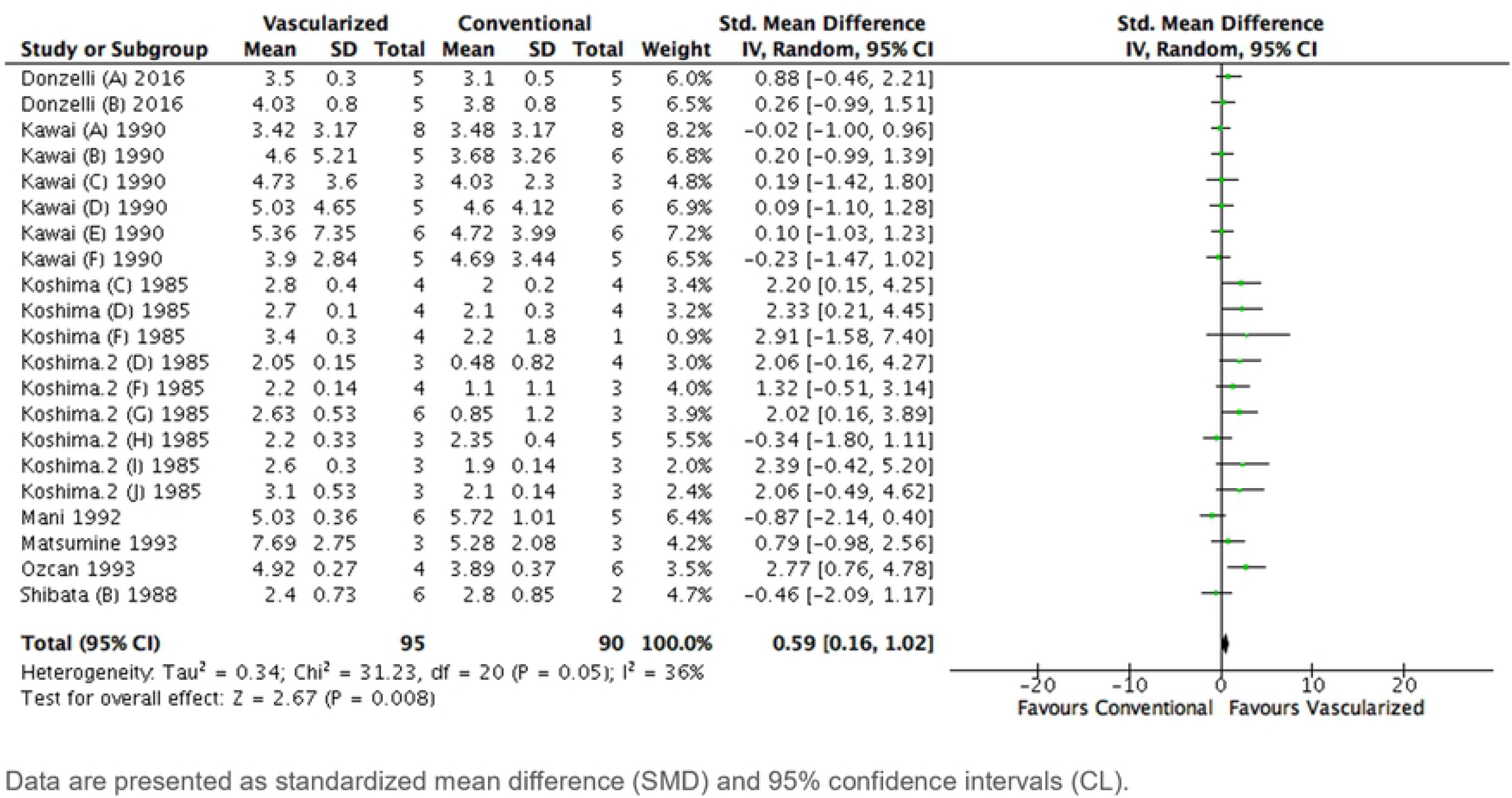
Forest plot of the effect of treatment with a vascularized n.

The subgroup analysis for graft length could not be interpreted because both the middle and long group consisted of fewer than 3 studies.

For species however, there was a significant difference in diameter comparing rabbits and rats showing a more positive result in rats (SEM 0.13 [95% CI -0.28 to 0.54], N = 11; I^2^ = 11% compared to SEM 1.40 [95% CI 0.74 to 2.06], N = 10; I^2^ = 3%; P = 0.005). Rats showed a significant larger nerve fiber diameter in vascularized grafts compared to conventional grafts (S4 Fig).

A significant difference in diameter could not be found comparing different time frames (S5 Fig).

### Nerve conduction velocity

Data on nerve conduction velocity could be extracted from 3 studies containing 4 comparisons.(21, 29, 31) Three comparisons used a rabbit model. A total of 74 animals were operated on, resulting in 74 grafts that met our selection criteria. Graft length ranged from 20 to 30 mm. Outcomes were measured at time points between 70 and 168 days.

Overall, analysis showed treatment with a vascularized nerve graft resulted in a significantly higher nerve conduction velocity (SMD, 1.19 [95% CI 0.19 to 2.19], N = 4) (Fig 6). Between studies, heterogeneity was high (I^2^ = 79%). There were not enough studies to perform a subgroup analysis.

**Fig 6.**
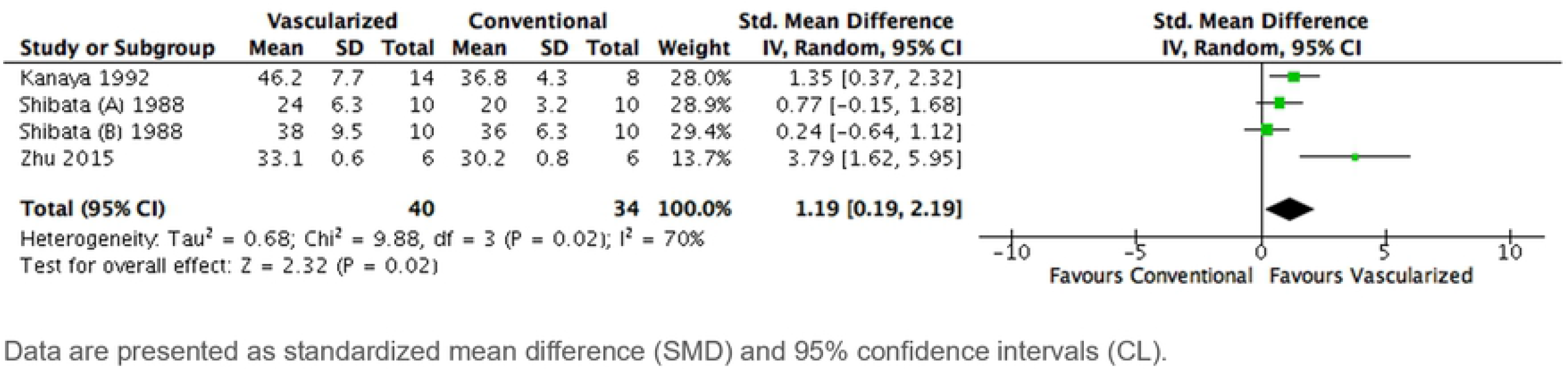
Forest plot of the effect of treatment with a vascularized n.

### Muscle weight

Two studies, containing 6 comparisons, assessed muscle weight.(18, 21) A total of 92 animals, all rats, were operated on, resulting in 92 grafts. The two graft lengths used were 20 and 25mm. The varying time points at which data were extracted were between 84 and 360 days.

Overall, no significant difference was found between the treatment groups (SMD, 0.18 [95% CI -0,24 to 0,60], N = 6), I^2^ was 0% (Fig 7). There were not enough studies to perform a subgroup analysis.

**Fig 7.**
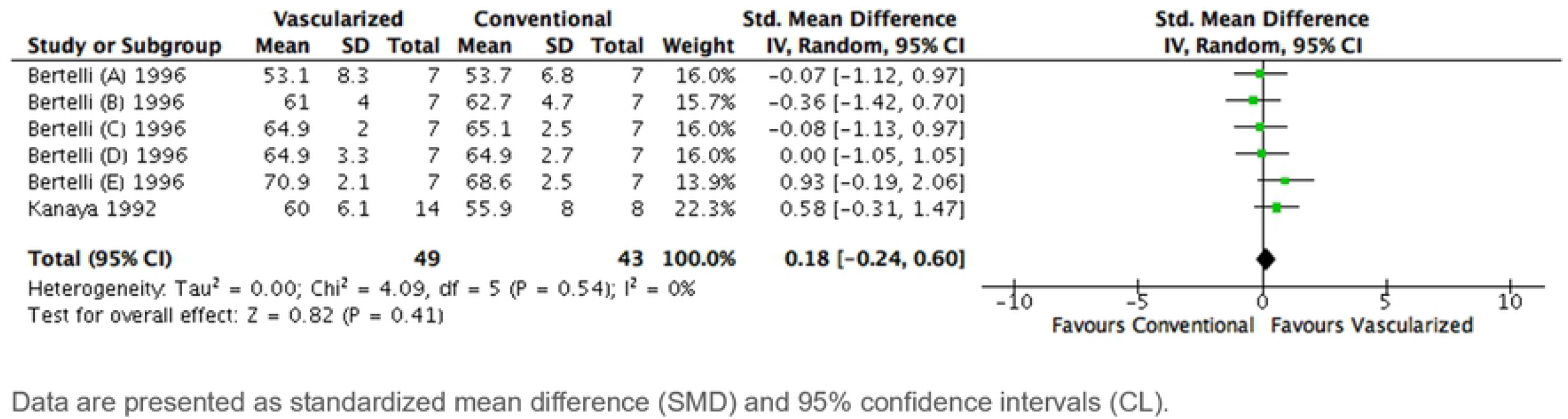
Forest plot of the effect of treatment with a vascularized n.

### Sensitivity analyses

#### Axonal count

Exclusion of the studies in which animals were their own control group altered our results significantly. The previous effect in favor of a vascularized nerve graft compared to a conventional nerve autograft was no longer available (SMD 0.26 [95% CL -0.09 to 0.62], N = 18), heterogeneity was I^2^ = 17% (Fig 8). Conclusions of all subgroup analyses appeared to be robust (S6 Fig, S7 Fig, S8 Fig)

**Fig 8.**
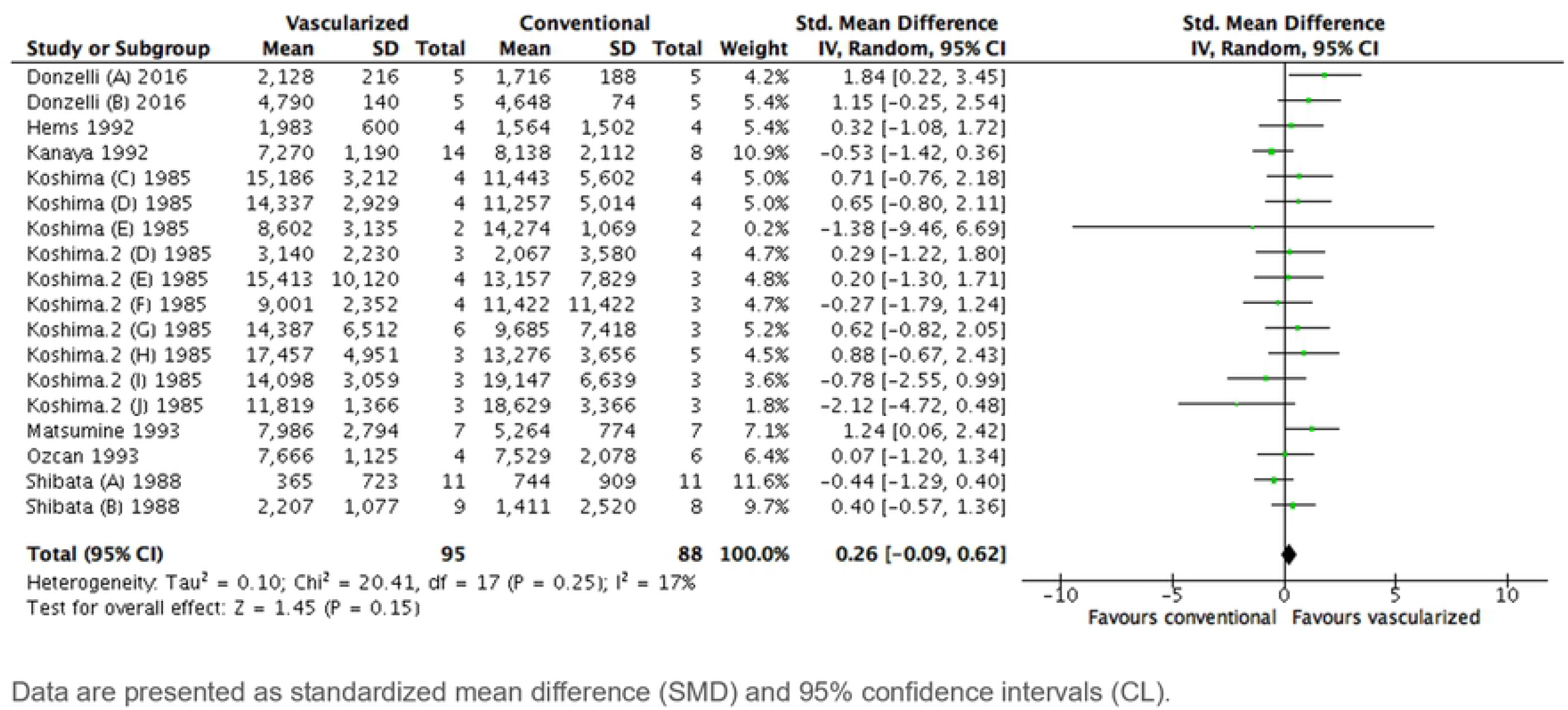
Sensitivity analysis, forest plot of the effect of treatment w.

#### Diameter

Exclusion of the studies in which animals were their own control group did not alter our results significantly. A significant difference in favor of a vascularized nerve graft compared to a conventional nerve autograft was found (SMD 1.03 [95% CL 0.39 to 1.68], N = 15), heterogeneity was I^2^ = 46% (Fig 9).

**Fig 9.**
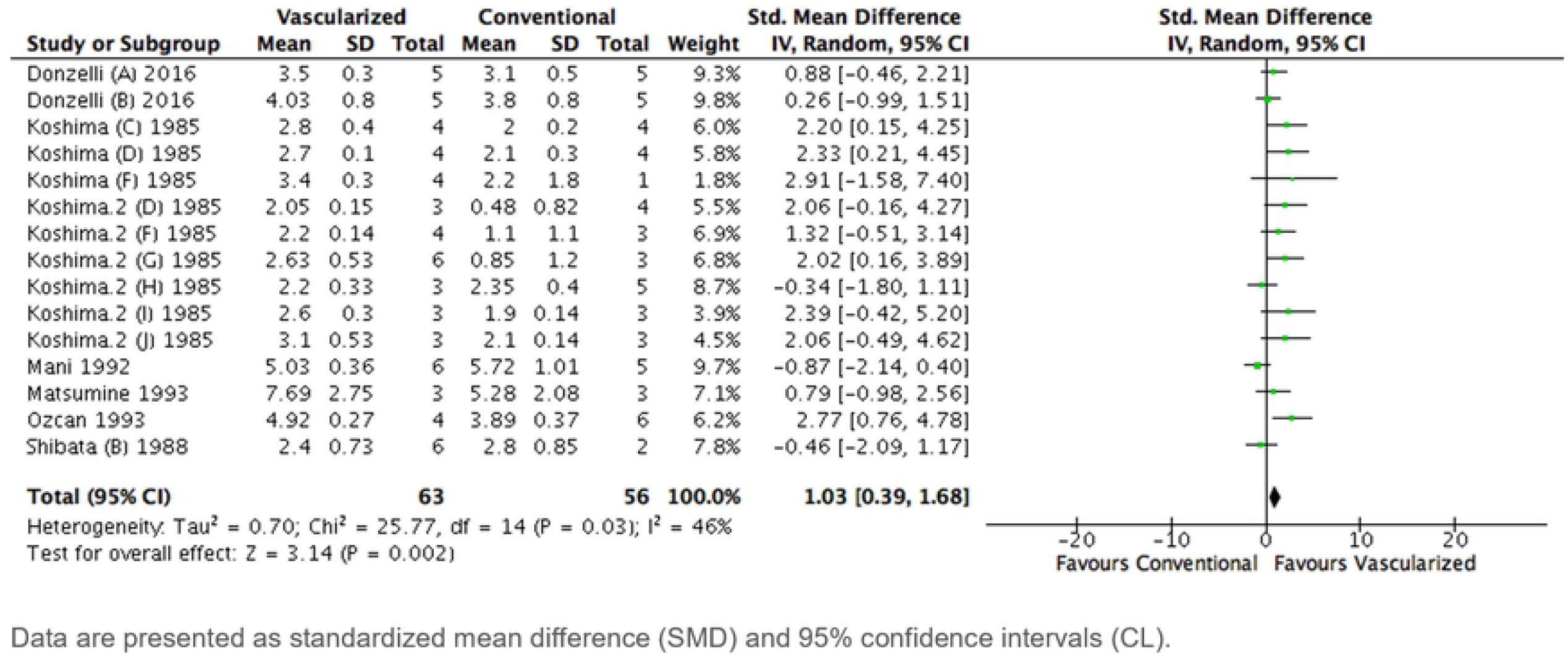
Sensitivity analysis, forest plot of the effect of treatment wi.

**Fig 10.**
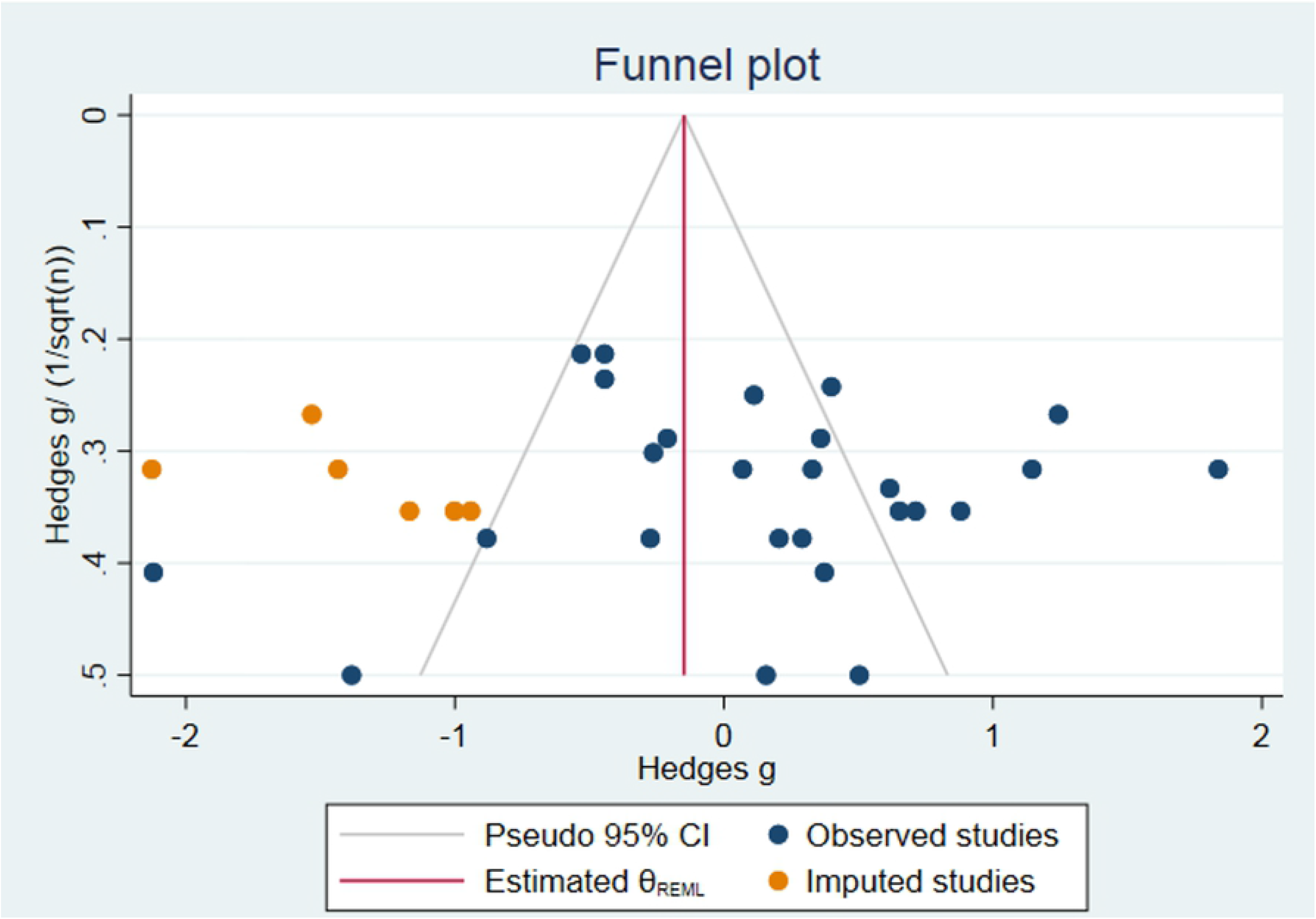
Publication bias.

Next to that, the result of the subgroup analysis on species was altered. No significant difference in favor of rats was found. (SEM 0.39 [95% CI -0.68 to 1.45], N = 5; I^2^ = 62% compared to SEM 1.40 [95% CI 0.74 to 2.06], N = 10; I^2^ = 3%; P = 0.13) (S9 Fig). Other conclusions appeared to be robust (S10 Fig)

### Publication bias analysis

Publication bias was assessed for axonal count only, because all other outcomes contained fewer than 10 studies.

#### Axonal Count

The funnel plot suggested some asymmetry. Duval and Tweedie’s Trim and Fill analysis resulted in 6 extra data points (see Fig 5), indicating the presence of publication bias and some overestimation of the identified summary effect size.

## Discussion

This review suggests that a vascularized nerve graft does result in a significantly better nerve recovery compared to non-vascularized nerve autografts in animal models regarding the outcome measurements nerve fiber diameter, nerve conduction velocity and axonal count. However, the effect on axonal count did not appear to be very robust as after sensitivity analysis the effect was no longer present. Muscle weight did not differ between vascularized and non-vascularized grafts. Subgroup analysis indicated that the effect of vascularized graft on nerve fiber diameter is larger in rats compared to rabbits. However, this difference disappeared after sensitivity analysis.

There is a lot of discussion on what the best outcome measurement for nerve regeneration is. Until this day there is no proper “gold standard” to test nerve recovery, although the ultimate goal of nerve recovery is to maximize sensation and motion. The most commonly used outcome measurement for sensation is the von Frey test.(32) For motion, walking track analysis was believed to be the best overall assessment.(33-36) At the moment it is rarely used and some would say it is even obsolete. Additionally, walking track analysis does not reflect maximum muscle force capacity. Others say the most precise measurement is the isometric response of muscle to tetanic contraction.(37) The authors are aware of the fact that histomorphometry, electrophysiology and axonal count in particular may be minimally correlated to the real functional recovery of sensation or motion.(38) Still, these were the outcome measurements used for want of better ones.

This present meta-analysis of animal studies is, to the best of our knowledge, the first of its kind. Only some human case reports exist to try to put our findings into a broader perspective.(8, 11, 39) The clinical observations in these human case reports did not include the outcome measurements of this review. Nevertheless, all showed a superior sensory recovery in vascularized nerve grafts compared to conventional nerve grafts using different outcome measurements, such as the presence of a sharp/blunt discrimination, cold intolerance, the Tinel’s sign and the Semmes-Weinstein monofilament test.

Notably, clinical case reports found that vascularized nerve grafts give a better recovery in large nerve grafts compared to conventional nerve autografts. Terzis et al. (40) showed that a vascularized nerve graft successfully bridges a nerve defect longer than 13 cm where conventional nerve grafts generally fail. Also Xu et al. (41) and Okinaga et al. (42) concluded that when the graft length was short, the results were not significantly in favor of a vascularized nerve graft. However, we did not find a difference in recovery between various graft lengths in this meta-analysis.

### Limitations of this review

Firstly, our risk of bias analysis showed that most studies reported poorly on important methodological details. Therefore, most of the risk of bias items assessed had to be scored as unclear risk of bias. Even though this is quite commonly seen in animal studies, it is something to be taken into account.(43) The absence of reporting such methodological details could, to a certain extent, indicate the negligence of using these methods to minimize bias and confounding.(44) This can seriously hamper the possibility to draw reliable conclusions from the included animal studies.

Secondly, the number of studies included in this meta-analysis is relatively low, especially on nerve conduction velocity and muscle weight. This resulted in subgroups being relatively small, even to the extent that some subgroup analysis could not be interpreted. Furthermore, heterogeneity was moderate to high. However, because of their explorative nature a moderate to high heterogeneity between animal studies is expected.

To account for anticipated heterogeneity, we used a random effects model, conducted sensitivity analyses and explored the suggested causes for between study heterogeneity by means of subgroup analyses. Exploring this heterogeneity is one of the added values of meta-analyses of animal studies and might help to inform the design of future animal studies and subsequent clinical trials.

Thirdly, the graft length used to repair a nerve defect in rat and rabbit models is presumably smaller than those needed in humans. Therefore, the results shown in these animal experiments might not be correlated with the expected clinical outcomes.

Fourthly, a possible reason for heterogeneity could be the use of animals as their own control in some studies. Therefore, a sensitivity analysis was performed. This led to 3 studies being excluded because animals were used as their own control group. When Kawai et al. (22), Seckel et al. (28) and Tark et al. (30) were excluded there was not a significant difference in axonal count in favor of vascularized nerve grafts compared to conventional nerve autografts.

Lastly, the presence of publication bias was identified. Our funnel plot suggested some asymmetry and Duval and Tweedie’s Trim and Fill analysis predicts some overestimation of the identified summary effect size of axonal count.

## Conclusion

Treating a nerve gap with a vascularized graft results in superior nerve recovery compared to non-vascularized autografts nerve grafts in three out of four outcome measurements.

However, this conclusion needs to be taken with some caution due to the inherent limitations of this meta-analysis. In addition, we recommend future studies to be performed under conditions more closely resembling human circumstances and to use long nerve grafts. Furthermore, we underline that future studies should use the Gold Standard Publication Checklist or ARRIVE guidelines to improve the reporting and methodological quality of animal studies.(45, 46) This is essential to improve the quality of the evidence presented in animal studies and the successful translation to humans in a clinical setting.

## Acknowledgements

The authors would like to thank Mrs On Ying Chan (Health Sciences reference librarian, Radboud University) for assisting with the development of the search strategy.

---

**S1 Fig.**
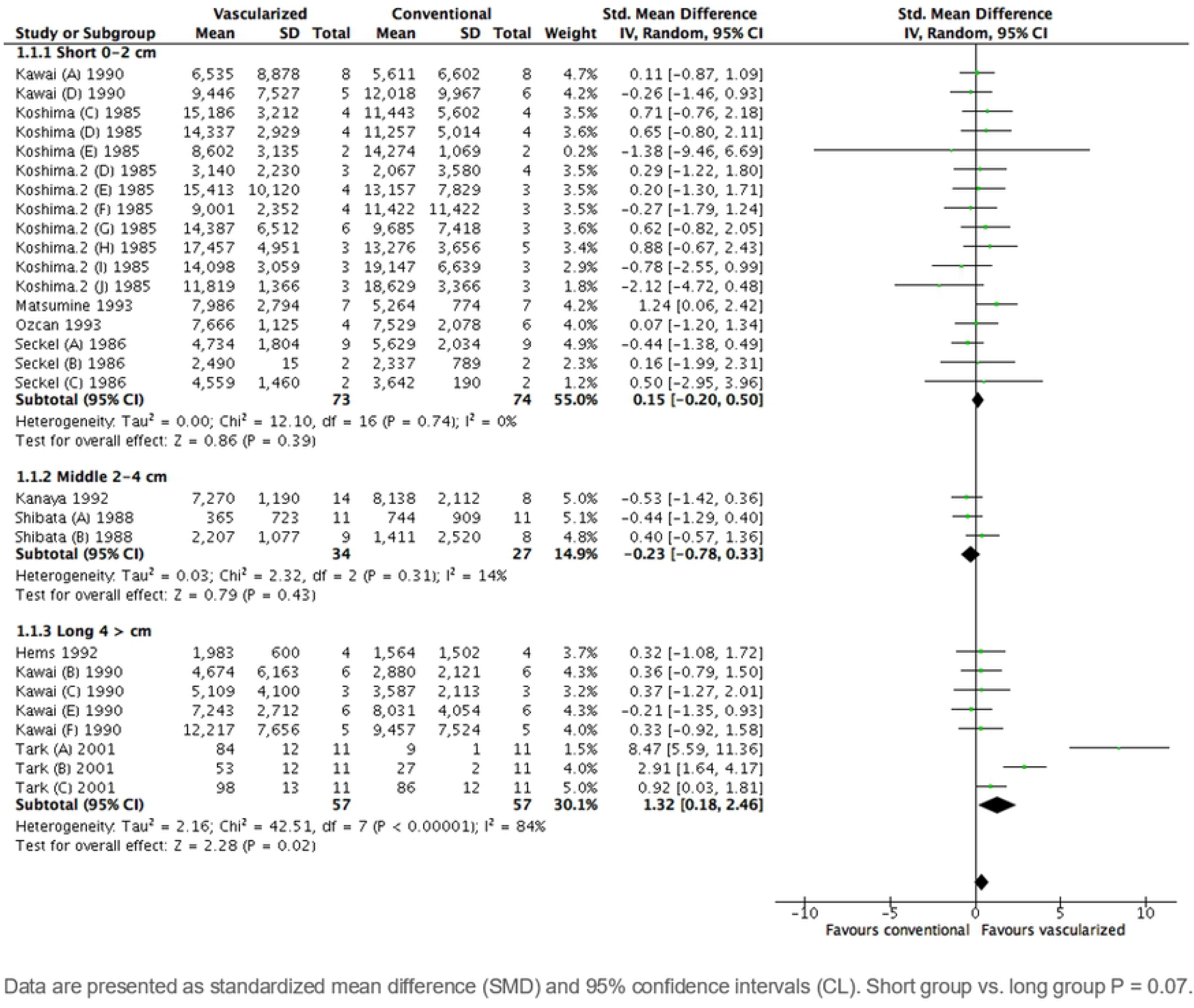
Subgroup analysis by graft length on axonal count.

**S2 Fig.**
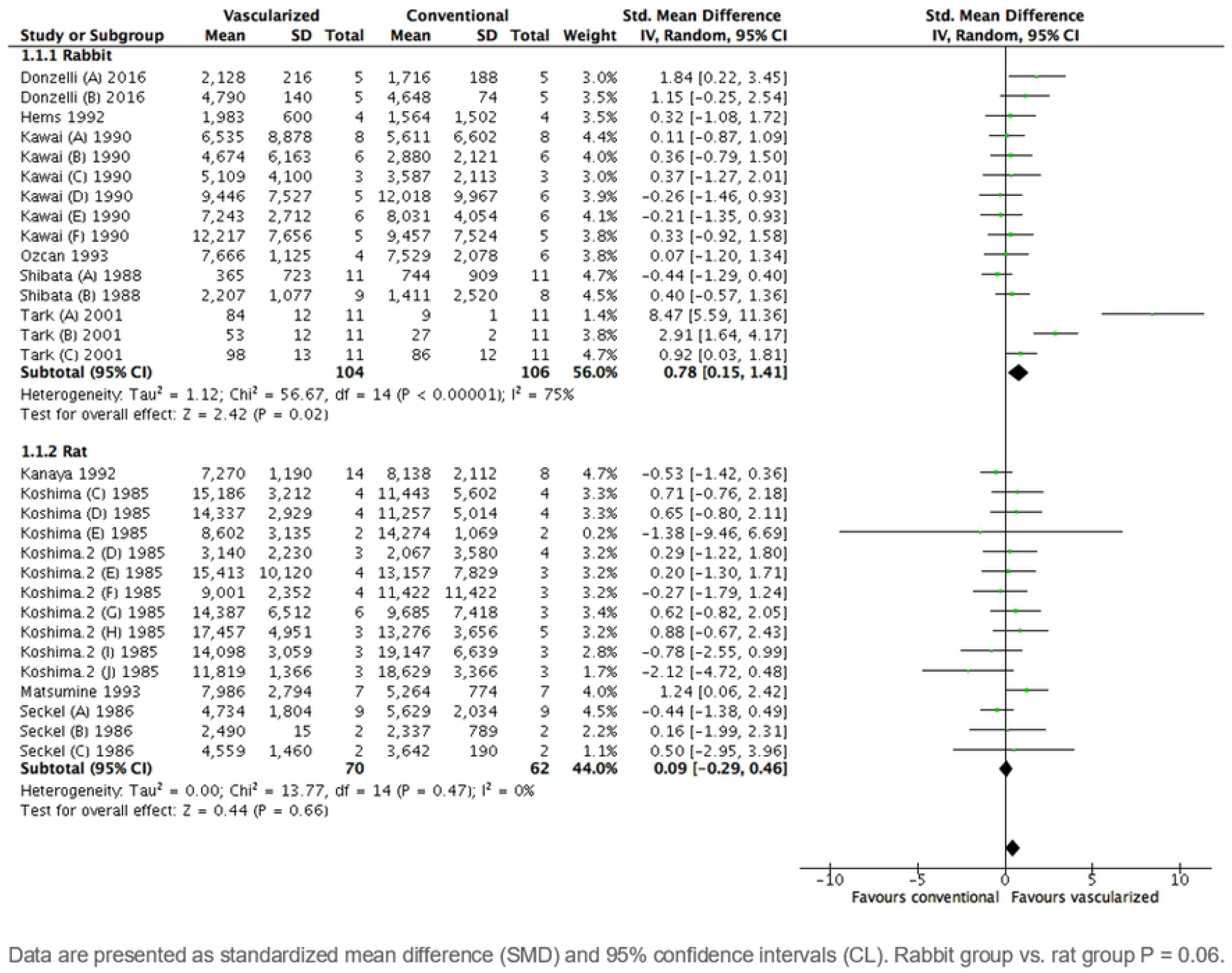
Subgroup analysis by graft length on axonal count.

**S3 Fig.**
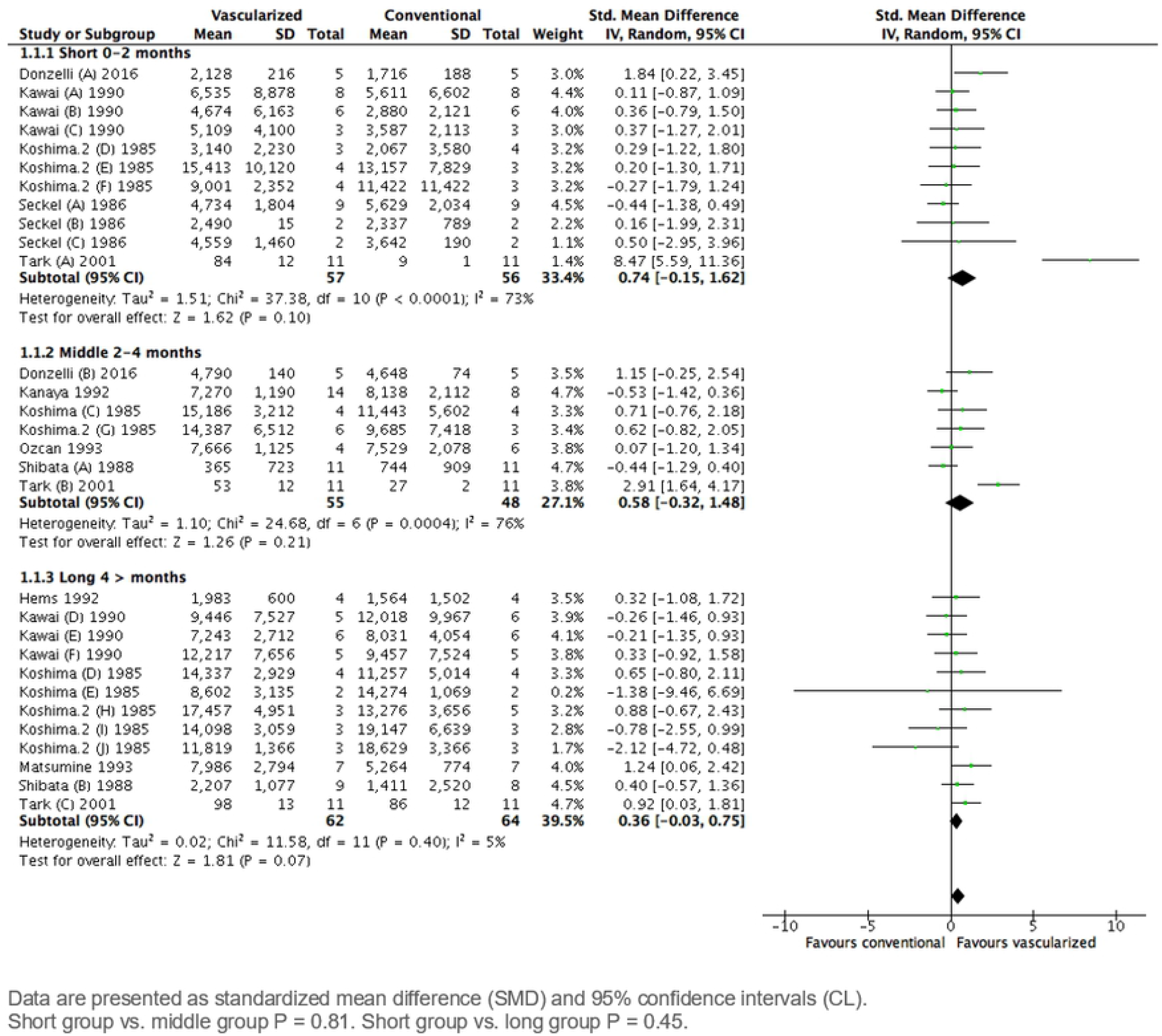
Subgroup analysis by graft length on axonal count.

**S4 Fig.**
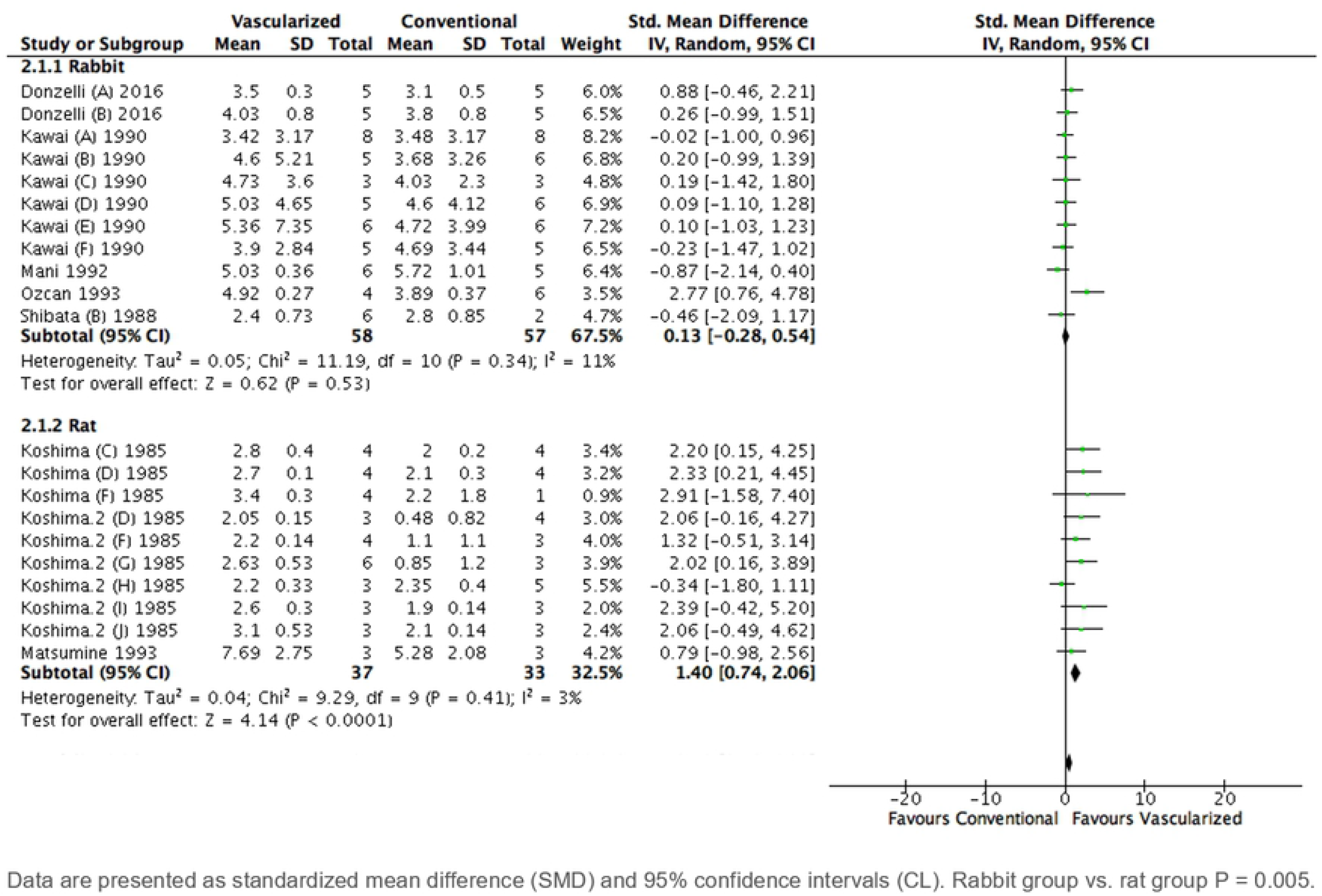
Subgroup analysis by species on diameter.

**S5 Fig.**
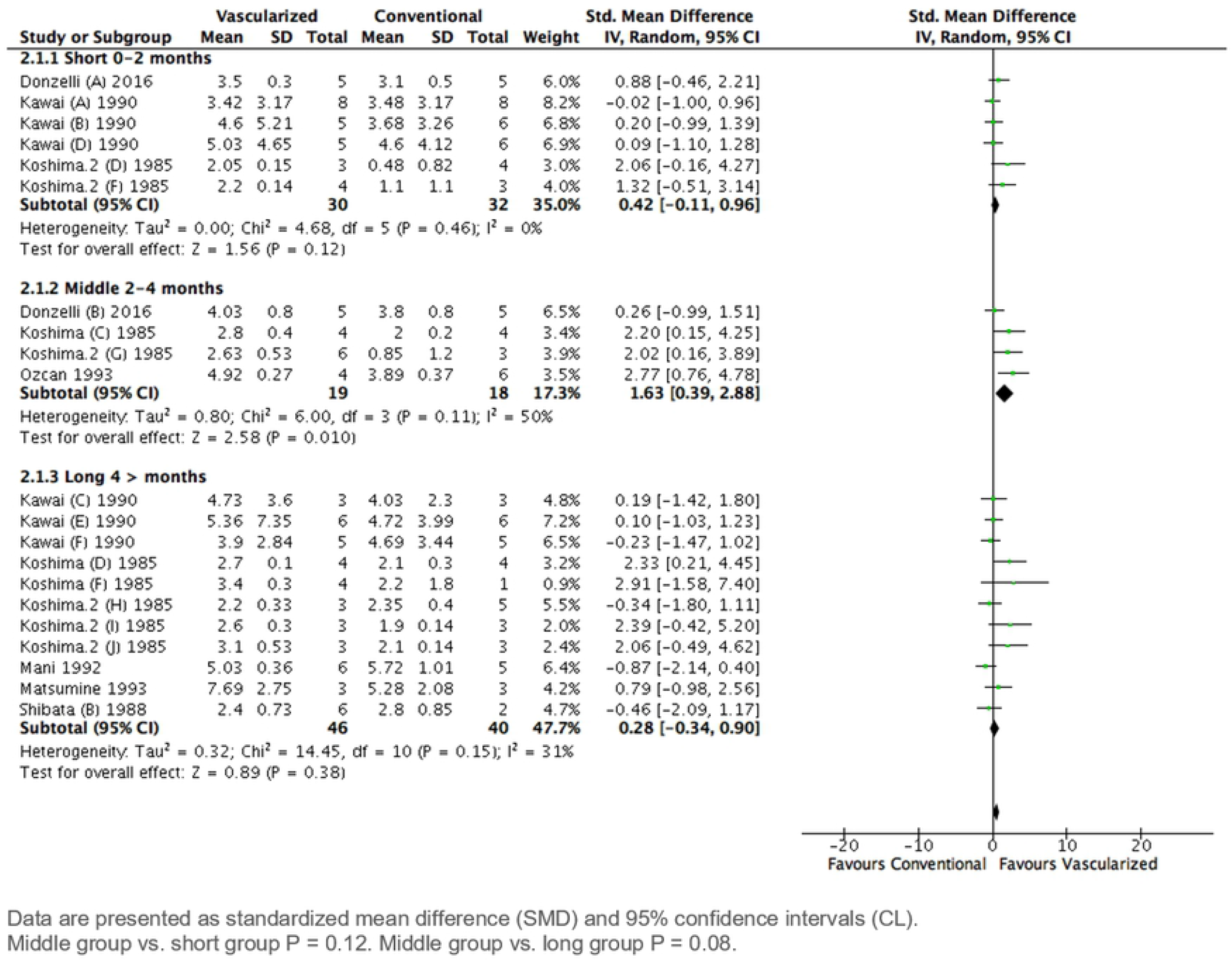
Subgroup analysis by time frame on diameter.

**S6 Fig.**
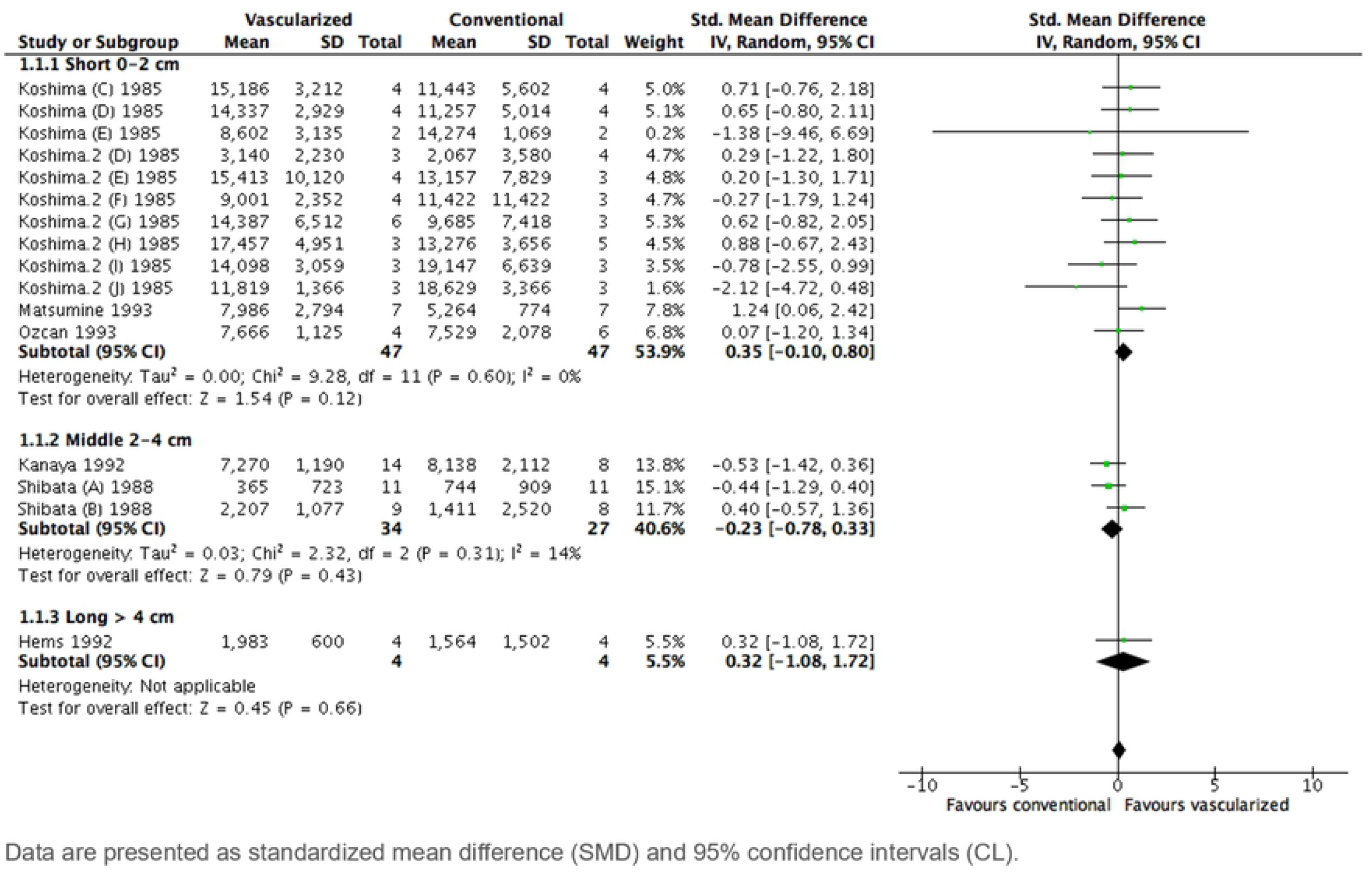
Sensitivity analysis subgroup by graft length on axonal count.

**S7 Fig.**
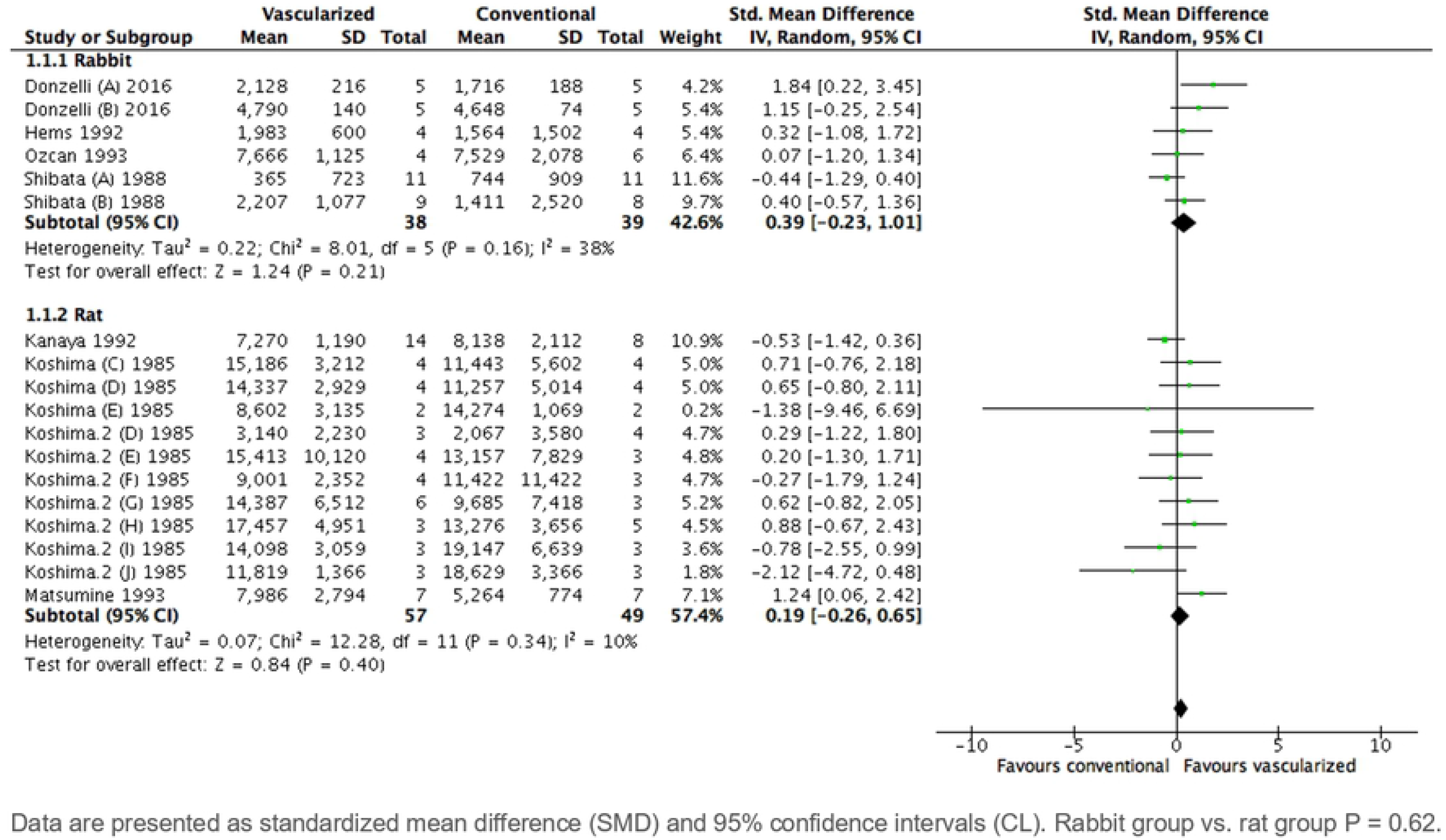
Sensitivity analysis subgroup by species on axonal count.

**S8 Fig.**
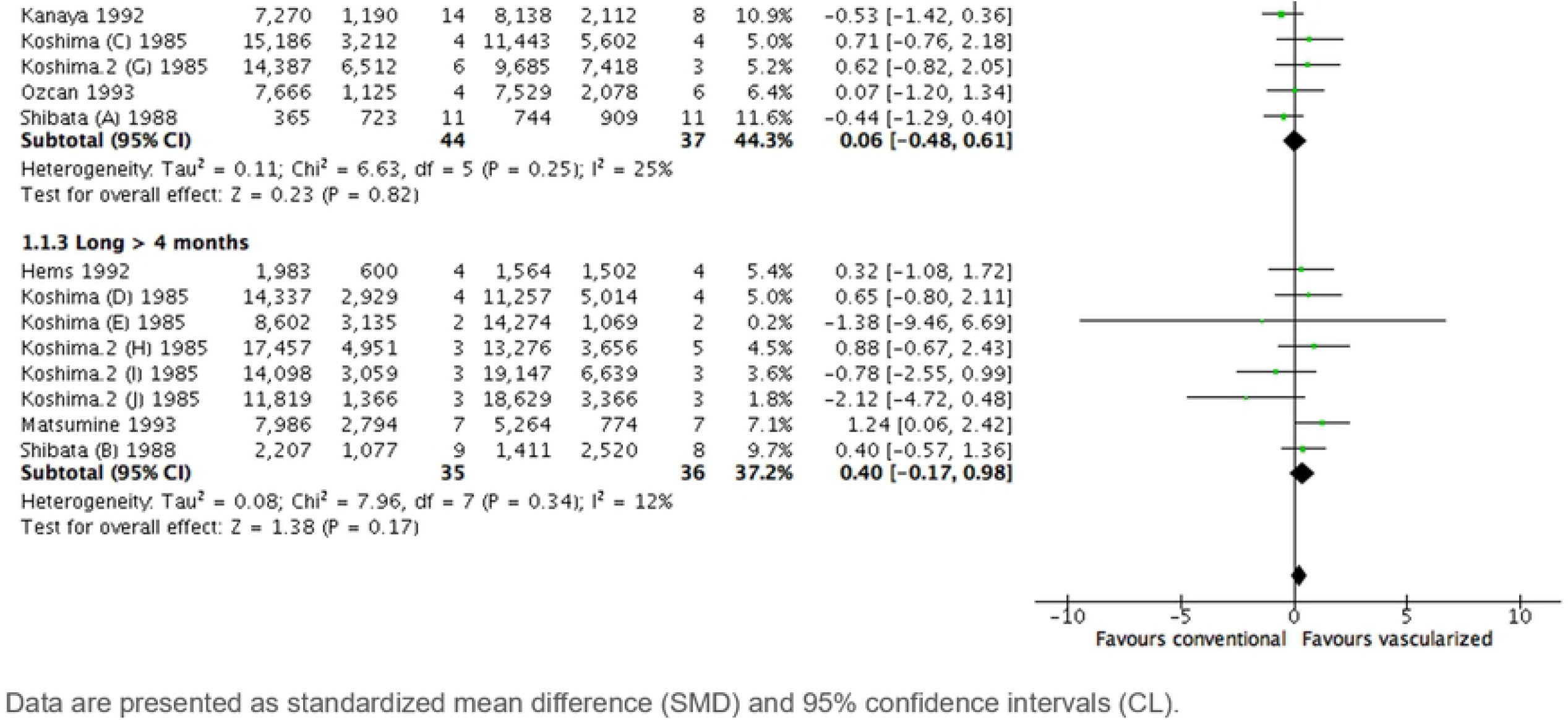
Sensitivity analysis subgroup by time frame on axonal count.

**S9 Fig.**
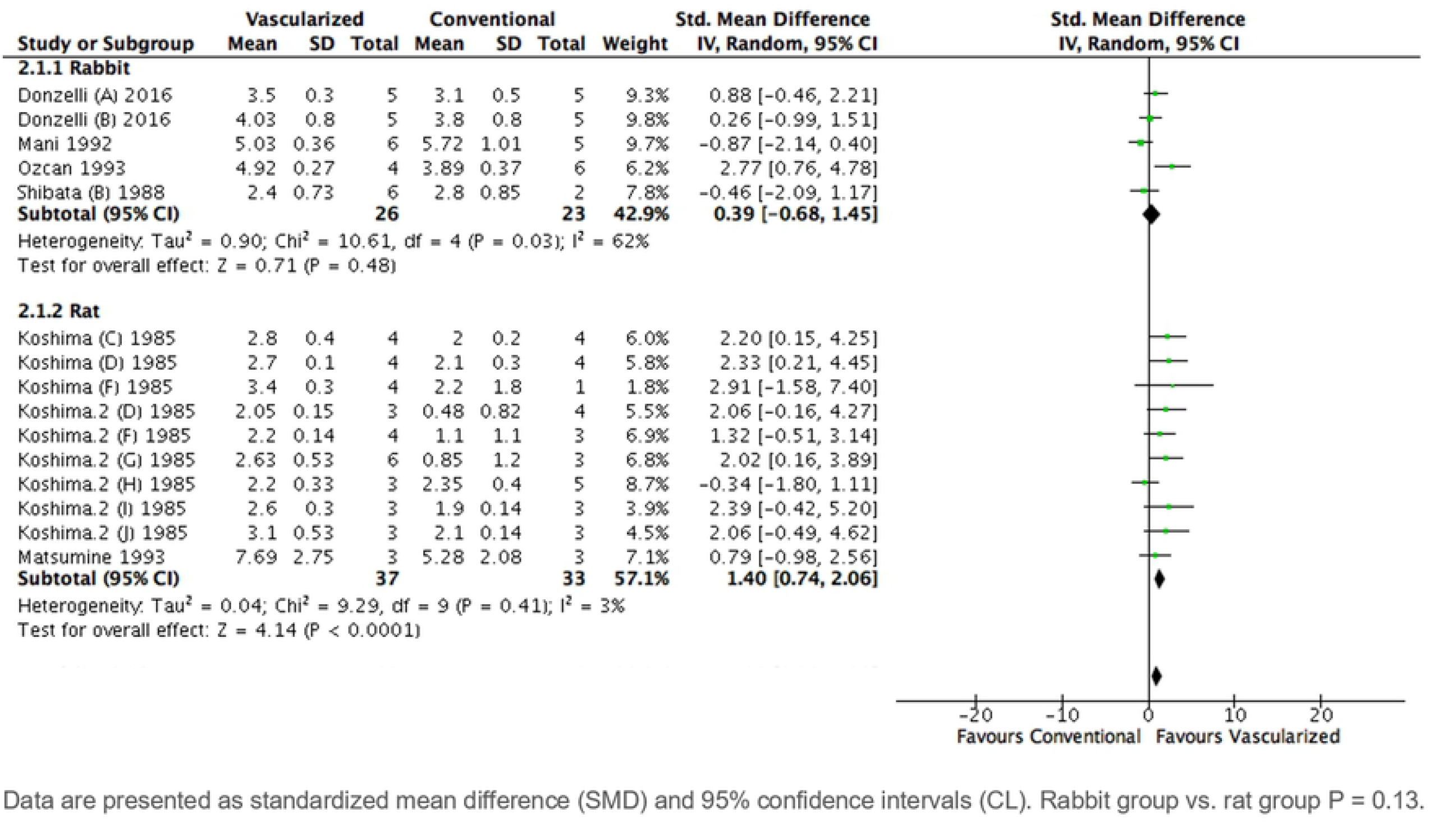
Sensitivity analysis subgroup by species on diameter.

**S10 Fig.**
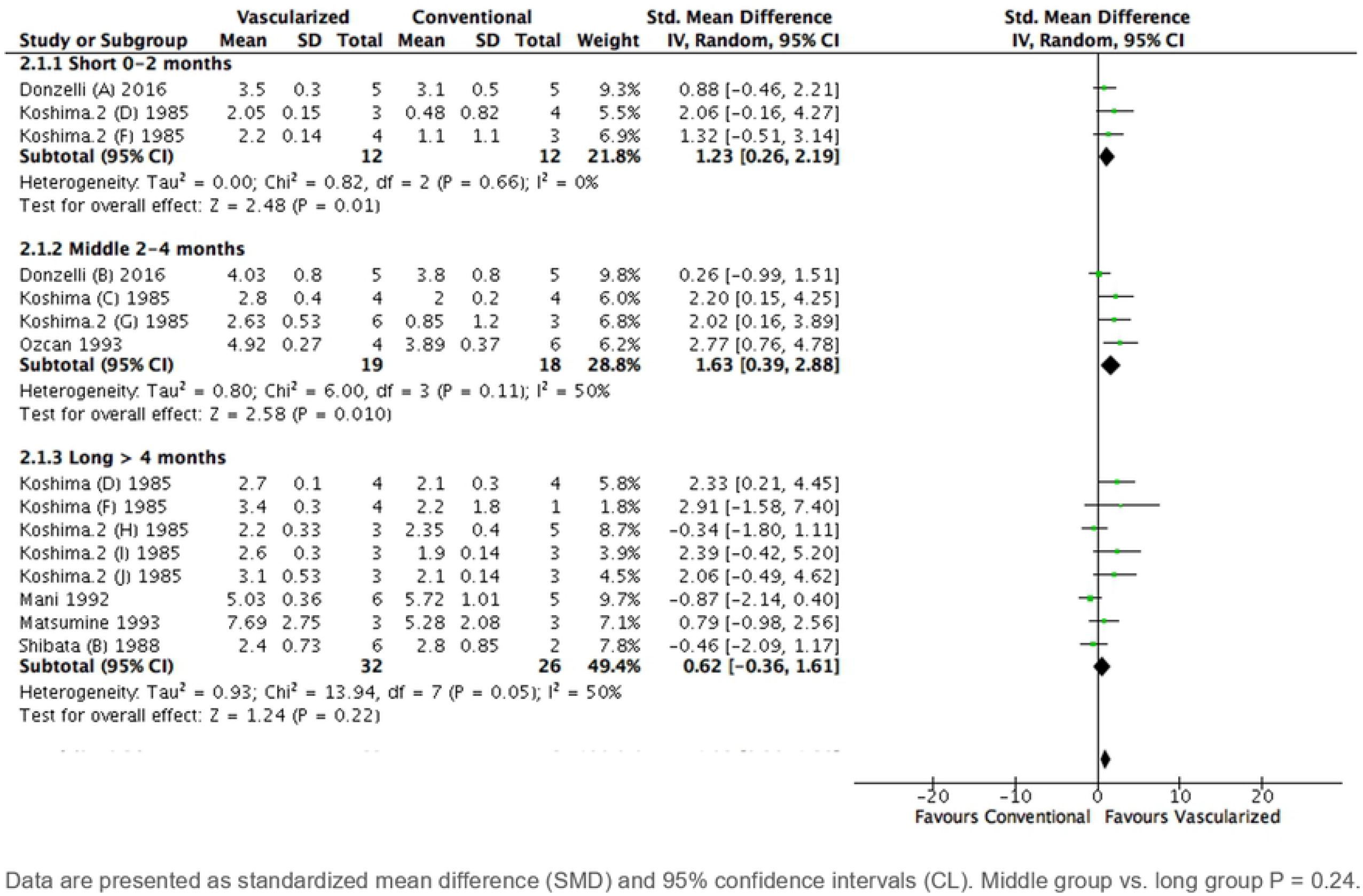
Sensitivity analysis subgroup by time frame on diameter.

**S1 Table.**
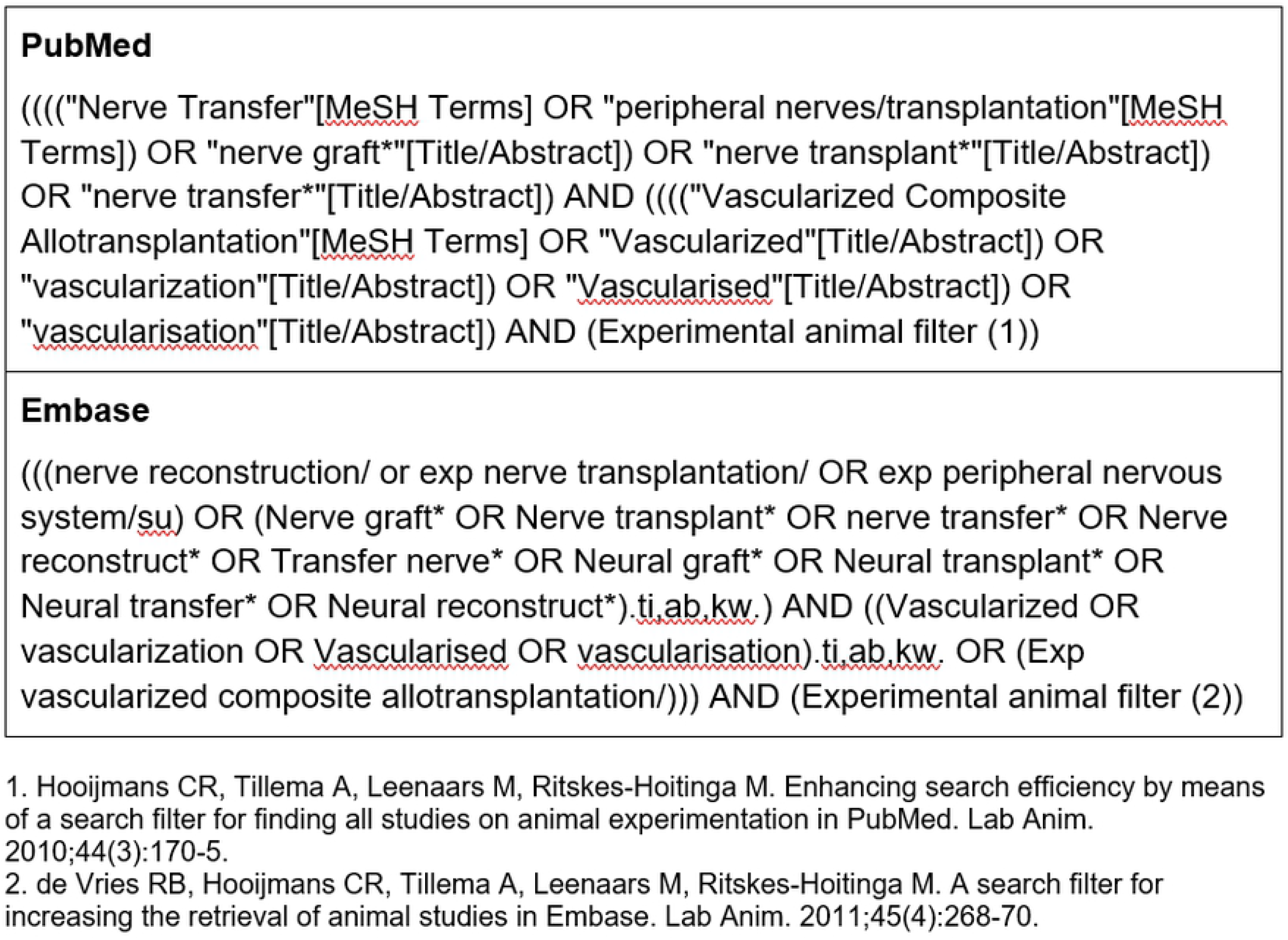
Search strategy

## References

1. Robinson PP, Loescher AR, Smith KG. A prospective, quantitative study on the clinical outcome of lingual nerve repair. Br J Oral Maxillofac Surg. 2000;38(4):255–63.

2. Gordin E, Lee TS, Ducic Y, Arnaoutakis D. Facial nerve trauma: evaluation and considerations in management. Craniomaxillofac Trauma Reconstr. 2015;8(1):1–13.

3. Seddon HJ. NERVE GRAFTING. J Bone Joint Surg Br. 1963;45:447–61.

4. Jain RK, Au P, Tam J, Duda DG, Fukumura D. Engineering vascularized tissue. Nat Biotechnol. 2005;23(7):821–3.

5. Tarlov IM EJ. Nerve grafts: the importance of an adequate blood supply. J Neurosurg. 1945;2:49.

6. Iida T, Nakagawa M, Asano T, Fukushima C, Tachi K. Free vascularized lateral femoral cutaneous nerve graft with anterolateral thigh flap for reconstruction of facial nerve defects. J Reconstr Microsurg. 2006;22(5):343–8.

7. Macionis V. A pedicled vascularized ulnar nerve graft based on the epineurial vascular supply: a case report. J Reconstr Microsurg. 2008;24(6):453–5.

8. Boorman JG, Sykes PJ. Vascularised versus conventional nerve grafting: a case report. J Hand Surg Br. 1987;12(2):218–20.

9. Halim AS, Yusof I. Composite vascularised osteocutaneous fibula and sural nerve graft for severe open tibial fracture--functional outcome at one year: a case report. J Orthop Surg (Hong Kong). 2004;12(1):110–3.

10. Luchetti R, De Santis G, Soragni O, Deluca S, Pederzini L, Alfarano M, et al. Vascularized nerve grafting: case reports. Rivista di Patologia dell’Apparato Locomotore. 1987;7(1-2):165–71.

11. Mackinnon SE, Kelly L, Hunter DA. Comparison of regeneration across a vascularized versus conventional nerve graft: case report. Microsurgery. 1988;9(4):226–34.

12. Usami S, Kawahara S, Inami K, Hirase Y. Use of a vascularized dorsal sensory branch of an ulnar nerve flap for repairing a proper digital nerve with coverage of a volar soft tissue defect: Report of two cases. Microsurgery. 2019;39(7):647–50.

13. Taylor GI, Ham FJ. The free vascularized nerve graft. A further experimental and clinical application of microvascular techniques. Plast Reconstr Surg. 1976;57(4):413–26.

14. Lux P, Breidenbach W, Firrell J, Wood MB. Determination of temporal changes in blood flow in vascularized and nonvascularized nerve grafts in the dog. Plastic and Reconstructive Surgery. 1988;82(1):133–44.

15. Hooijmans CR, Tillema A, Leenaars M, Ritskes-Hoitinga M. Enhancing search efficiency by means of a search filter for finding all studies on animal experimentation in PubMed. Lab Anim. 2010;44(3):170–5.

16. de Vries RB, Hooijmans CR, Tillema A, Leenaars M, Ritskes-Hoitinga M. A search filter for increasing the retrieval of animal studies in Embase. Lab Anim. 2011;45(4):268–70.

17. Hooijmans CR, Rovers MM, de Vries RB, Leenaars M, Ritskes-Hoitinga M, Langendam MW. SYRCLE’s risk of bias tool for animal studies. BMC Med Res Methodol. 2014;14:43.

18. Bertelli JA, Taleb M, Mira JC, Calixto JB. Muscle fiber type reorganization and behavioral functional recovery of rat median nerve repair with vascularized or conventional nerve grafts. Restor Neurol Neurosci. 1996;10(1):5–12.

19. Donzelli R, Capone C, Sgulò FG, Mariniello G, Maiuri F. Vascularized nerve grafts: an experimental study. Neurol Res. 2016;38(8):669–77.

20. Hems TEJ, Glasby MA. Comparison of different methods of repair of long peripheral nerve defects: An experimental study. British Journal of Plastic Surgery. 1992;45(7):497–502.

21. Kanaya F, Firrell J, Tsai TM, Breidenbach WC. Functional results of vascularized versus nonvascularized nerve grafting. Plast Reconstr Surg. 1992;89(5):924–30.

22. Kawai H, Baudrimont M, Travers V, Sedel L. A comparative experimental study of vascularized and nonvascularized nerve grafts. J Reconstr Microsurg. 1990;6(3):255–9.

23. Koshima I, Harii K. Experimental study of vascularized nerve grafts: morphometric study of axonal regeneration of nerves transplanted into silicone tubes. Ann Plast Surg. 1985;14(3):235–43.

24. Koshima I, Harii K. Experimental study of vascularized nerve grafts: multifactorial analyses of axonal regeneration of nerves transplanted into an acute burn wound. J Hand Surg Am. 1985;10(1):64–72.

25. Mani GV, Shurey C, Green CJ. Is early vascularization of nerve grafts necessary? J Hand Surg Br. 1992;17(5):536–43.

26. Matsumine H, Sasaki R, Takeuchi Y, Miyata M, Yamato M, Okano T, et al. Vascularized versus nonvascularized island median nerve grafts in the facial nerve regeneration and functional recovery of rats for facial nerve reconstruction study. J Reconstr Microsurg. 2014;30(2):127–36.

27. Ozcan G, Shenaq S, Mirabi B, Spira M. Nerve regeneration in a bony bed: vascularized versus nonvascularized nerve grafts. Plast Reconstr Surg. 1993;91(7):1322–31.

28. Seckel BR, Ryan SE, Simons JE, Gagne RG, Watkins E, Jr. Vascularized versus nonvascularized nerve grafts: an experimental structural comparison. Plast Reconstr Surg. 1986;78(2):211–20.

29. Shibata M, Tsai TM, Firrell J, Breidenbach WC. Experimental comparison of vascularized and nonvascularized nerve grafting. J Hand Surg Am. 1988;13(3):358–65.

30. Tark KC, Roh TS. Morphometric study of regeneration through vascularized nerve graft in a rabbit sciatic nerve model. J Reconstr Microsurg. 2001;17(2):109–14.

31. Zhu Y, Liu S, Zhou S, Yu Z, Tian Z, Zhang C, et al. Vascularized versus nonvascularized facial nerve grafts using a new rabbit model. Plast Reconstr Surg. 2015;135(2):331e–9e.

32. Kemp SW, Cederna PS, Midha R. Comparative outcome measures in peripheral regeneration studies. Exp Neurol. 2017;287(Pt 3):348–57.

33. Dellon AL, Mackinnon SE. Selection of the appropriate parameter to measure neural regeneration. Ann Plast Surg. 1989;23(3):197–202.

34. de Medinaceli L. Interpreting nerve morphometry data after experimental traumatic lesions. J Neurosci Methods. 1995;58(1-2):29–37.

35. Hadlock TA, Koka R, Vacanti JP, Cheney ML. A comparison of assessments of functional recovery in the rat. J Peripher Nerv Syst. 1999;4(3-4):258–64.

36. Koka R, Hadlock TA. Quantification of functional recovery following rat sciatic nerve transection. Exp Neurol. 2001;168(1):192–5.

37. Frykman GK, McMillan PJ, Yegge S. A review of experimental methods measuring peripheral nerve regeneration in animals. Orthop Clin North Am. 1988;19(1):209–19.

38. Wilbourn AJ. The electrodiagnostic examination with peripheral nerve injuries. Clin Plast Surg. 2003;30(2):139–54.

39. Chen C, Tang P, Zhang X. Reconstruction of proper digital nerve defects in the thumb using a pedicle nerve graft. Plast Reconstr Surg. 2012;130(5):1089–97.

40. Terzis JK, Kostopoulos VK. Vascularized nerve grafts for lower extremity nerve reconstruction. Ann Plast Surg. 2010;64(2):169–76.

41. Xu WD, Xu JG, Gu YD. Comparative clinic study on vascularized and nonvascularized full-length phrenic nerve transfer. Microsurgery. 2005;25(1):16–20.

42. Okinaga S, Nagano A. Can vascularization improve the surgical outcome of the intercostal nerve transfer for traumatic brachial plexus palsy? A clinical comparison of vascularized and non-vascularized methods. Microsurgery. 1999;19(4):176–80.

43. Macleod MR, van der Worp HB, Sena ES, Howells DW, Dirnagl U, Donnan GA. Evidence for the efficacy of NXY-059 in experimental focal cerebral ischaemia is confounded by study quality. Stroke. 2008;39(10):2824–9.

44. Hirst JA, Howick J, Aronson JK, Roberts N, Perera R, Koshiaris C, et al. The need for randomization in animal trials: an overview of systematic reviews. PLoS One. 2014;9(6):e98856.

45. Hooijmans CR, Leenaars M, Ritskes-Hoitinga M. A gold standard publication checklist to improve the quality of animal studies, to fully integrate the Three Rs, and to make systematic reviews more feasible. Altern Lab Anim. 2010;38(2):167–82.

46. Percie du Sert N, Hurst V, Ahluwalia A, Alam S, Avey MT, Baker M, et al. The ARRIVE guidelines 2.0: Updated guidelines for reporting animal research. PLoS Biol. 2020;18(7):e3000410.

